# Objects seen as scenes: neural circuitry for attending whole or parts

**DOI:** 10.1101/771899

**Authors:** Mitchell Valdés-Sosa, Marlis Ontivero-Ortega, Jorge Iglesias-Fuster, Agustin Lage-Castellanos, Jinnan Gong, Cheng Luo, Ana Maria Castro-Laguardia, Maria Antonieta Bobes, Daniele Marinazzo, Dezhong Yao

## Abstract

Depending on our goals, we pay attention to the global shape of an object or to the local shape of its parts, since it’s difficult to do both at once. This typically effortless process can be impaired in disease. However, it is not clear which cortical regions carry the information needed to constrain shape processing to a chosen global/local level. Here, novel stimuli were used to dissociate functional MRI responses to global and local shapes. This allowed identification of cortical regions containing information about level (independent from shape). Crucially, these regions overlapped part of the cortical network implicated in scene processing. As expected, shape information (independent of level) was mainly located in category-selective areas specialized for object- and face-processing. Regions with the same informational profile were strongly linked (as measured by functional connectivity), but were weak when the profiles diverged. Specifically, in the ventral-temporal-cortex (VTC) regions favoring level and shape were consistently separated by the mid-fusiform sulcus (MFS). These regions also had limited crosstalk despite their spatial proximity, thus defining two functional pathways within VTC. We hypothesize that object hierarchical level is processed by neural circuitry that also analyses spatial layout in scenes, contributing to the control of the spatial-scale used for shape recognition. Use of level information tolerant to shape changes could guide whole/part attentional selection but facilitate illusory shape/level conjunctions under impoverished vision.

**Significance statement:** One daily engages hierarchically organized objects (e.g. face-eyes-eyelashes). Their perception is commonly studied with global shapes composed by of local shapes. Seeing shape at one level is easy, but difficult for both at once. How can the brain guide attention to one level? Here using novel stimuli that dissociate different levels over time and examining local patterns of brain-activity, we found that the level and shape of visual objects were represented into segregated sets of cortical regions, each connected into their own pathway. Level information was found in part of the cortical network known to process scenes. Coding of object-level independently from shape could participate in guiding sustained attention within objects, eliminating interference from irrelevant levels. It could also help produce “illusory conjunctions” (perceptual migration of a shape to the wrong level) when attention is limited.

**Highlights:**

- Modified Navon figures allow dissociation in time of fMRI responses for the global/local levels.
- Shape-invariant hierarchical level information was found in scenes selective areas, whereas level-invariant shape information was found in object- and faces- selective areas.
- Level and shape regions were divided by the mid-fusiform sulcus (MFS) in VTC cortex, and each type of region connected into its own pathway.
- Having separate level/shape pathways could facilitate selective-attention, but foster illusory conjunctions.

## Introduction

Our visual world is full of hierarchically organized objects, and at different times it is more important to attend one echelon than the others (Kimchi, 2015). Consider identifying either the overall shape of a tree (a whole) in contrast to identifying the form of a leaf (a part). To orient attention according to hierarchical-level (i.e. global/local) it must be represented in the cerebral cortex, independently from shape and other visual attributes. But where and how? Furthermore, if cortical patches specialized in representing hierarchically-level do exist, into which neural circuits do they wire? For object shape, these questions have clear answers. Visual shape is recognized along a series of strongly connected cortical patches within the lateral ventral temporal cortex (VTC) (Grill-Spector and Weiner, 2014; Moeller et al., 2008), where increased tolerance to features not essential for object recognition (e.g. size, position, and viewpoint), as well as larger receptive fields (RFs), emerge by stages. It is also known that shape information reliably maps to category-specific cortical areas (Kanwisher, 2010).

In contrast, the cortical mapping of hierarchical-level information in itself (i.e. tolerant to variations in other visual properties) is unclear. On one hand, hierarchical-level (henceforth level) and shape are coded conjointly in early visual areas, where rudimentary attention to wholes and parts could arise by respectively selecting larger or smaller regions of visuotopic cortex, that is by varying the attentional-zoom (Sasaki et al., 2001). This means concentrating attention more towards the fovea for local shapes, but expanding it more peripherally for global shapes. On the other hand, for higher-order visual areas we can posit alternative hypothesis about shape-invariant level information. Firstly (H1), features associated with level (i.e. object-size, but also more complex attributes) could simply be discarded during extraction of level-invariant representations of shape in VTC (Rust and DiCarlo, 2010). Another hypothesis (H2), suggested by research in monkey VTC (Hong et al., 2016), is that codes for shape/level (invariant to each other) emerge together in the same pathway. Finally (H3), invariant codes for level (ignoring shape) could be extracted within a yet unidentified pathway, parallel to the route extracting shape within or outside of VTC. This would be analogous to a circuit in monkey VTC (Chang et al., 2017), in which information about color hue is refined across cortical patches while of information about shape is reduced. Since the input and output connections of any cortical region determine its function (Saygin et al., 2016), these hypothesis imply also different functional wiring schemes (Osher et al., 2016).

The aims of this article can be encapsulated in three questions (see Figure 1A): The first, where can we find shape-invariant level (and level-invariant shape) information in the cortex and which of the three hypothesis (H1-H3) is the most valid? Second, what is the relationship between the identified sites and previously characterized visual areas? Third, are the informational specialization and the functional connectivity of cortical patches related? To answer these questions we combined the use of novel stimuli with multivariate pattern and connectivity analyses of functional magnetic resonance (fMRI) activity.

**Figure 1.**
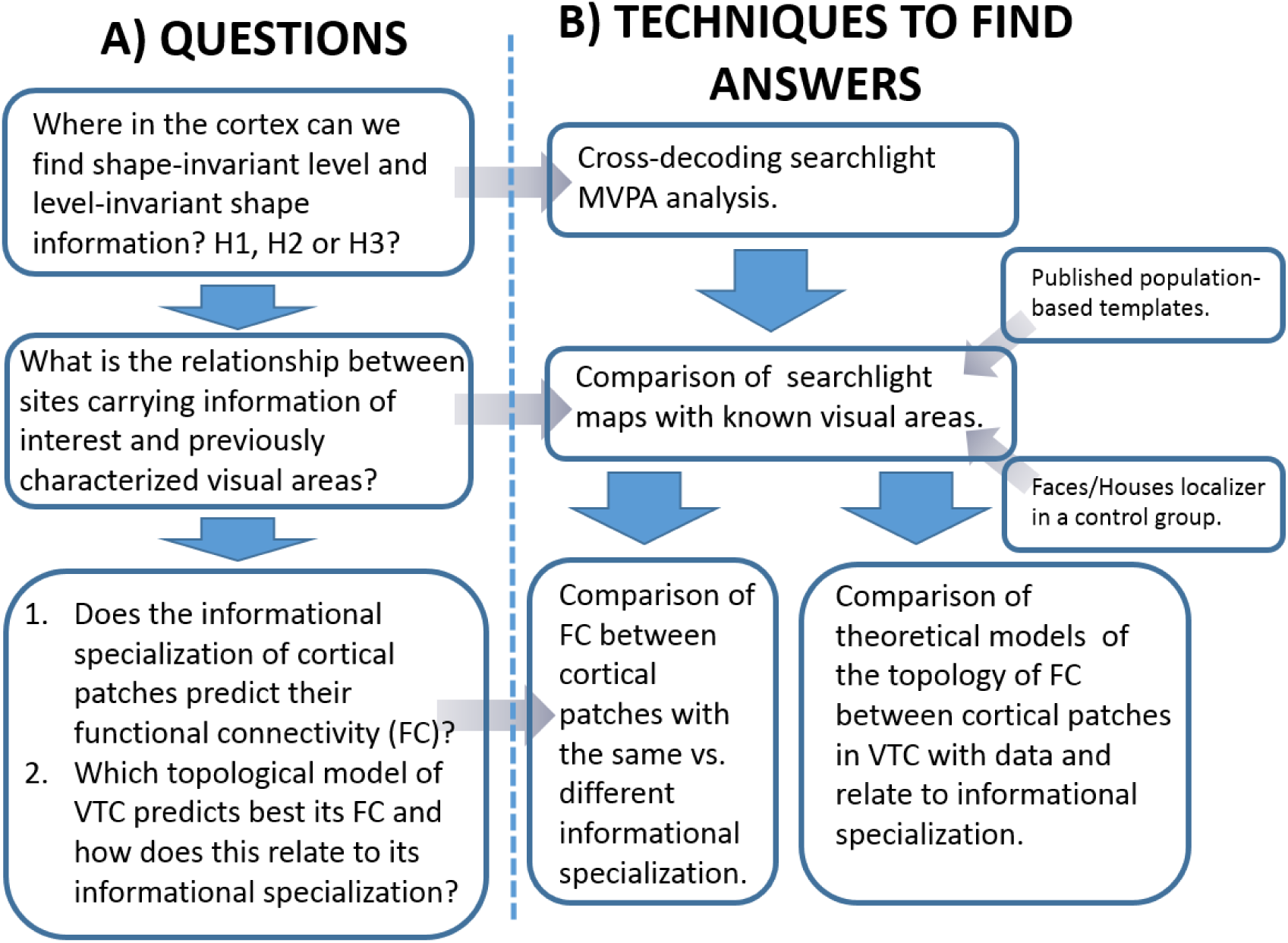
Flowchart outlining relationship between question and methods A) Questions presented in the introduction, where the alternative hypothesis H1 to H3 are explained. B) Overview of the methods used to answer the questions and their interrelations.

Experiments on the cognitive/neural mechanisms of level processing have traditionally used Navon figures (Fig. 2A): global shapes made out of local shapes (Navon, 2003). In real-world examples, shapes, cognitive tasks, and affordances usually differ across the global and local levels, but these can be equated between levels with these stimuli. Hence, differences in behavioral or neural responses to global vs. local shapes would largely depend on perceptual/attentional factors. However, one limitation of Navon figures (of particular relevance for our study) is that neural responses elicited by each level are difficult to separate for analysis. This is a consequence of the fact that global and local shapes onset/offset at the same time To solve this problem, we developed modified-Navon figures (Fig. 2B) (available at https://github.com/globallocal2019/Neuroimage-Paper-2019) that make it possible to present the global and the local elements dissociated over time (López et al., 2002). This provides the leverage needed here to analytically separate the neural responses elicited by each level.

**Figure 2.**
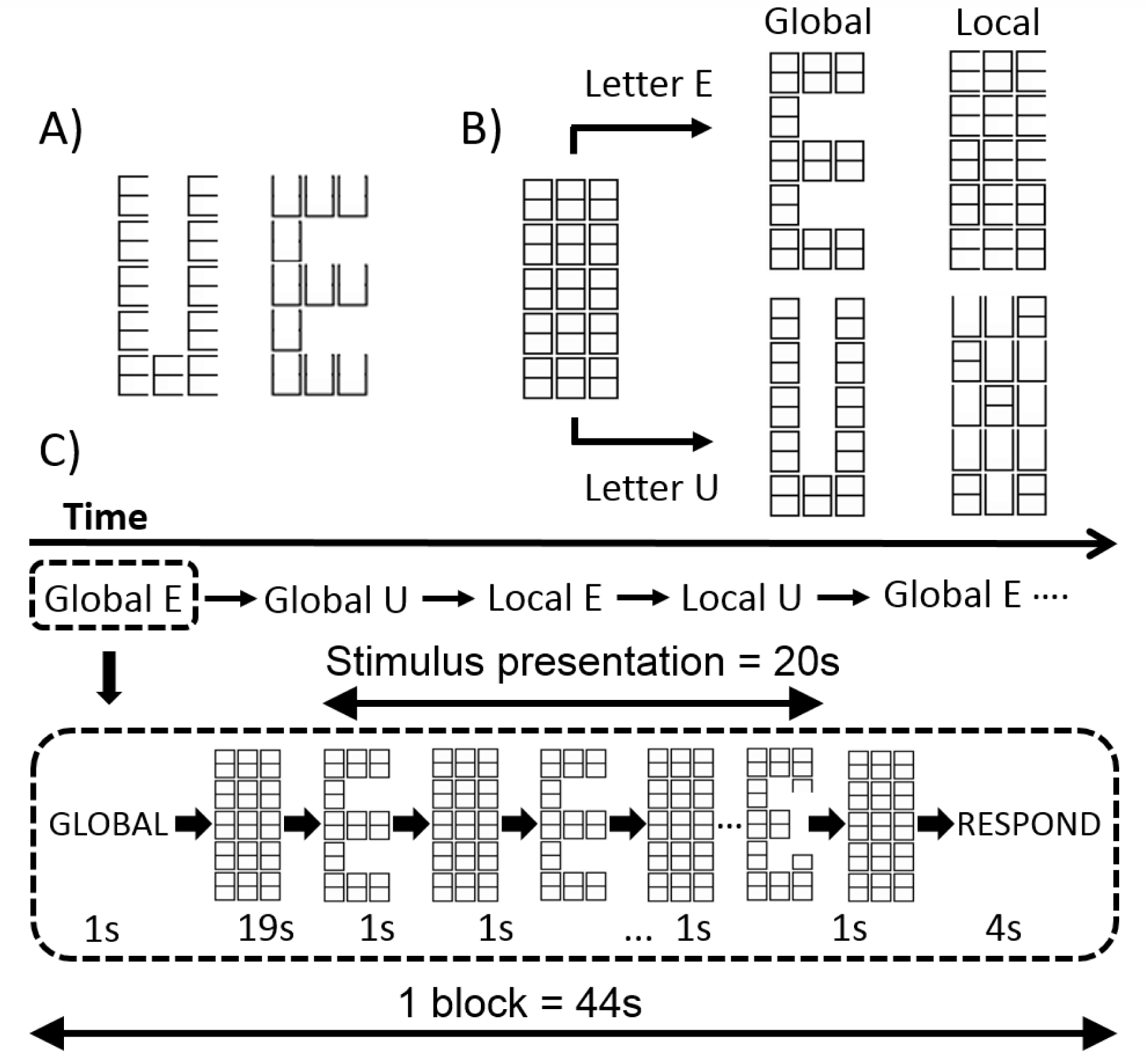
Stimuli and experimental design. A) Two traditional Navon figures: a global ‘U’ made of local ‘E’s, and a global ‘E’ made of local ‘U’s. Note that letters are present simultaneously at both levels. B) Modified Navon figures which emerged out of a mask. Two letter shapes (‘E’ and ‘U’) and two levels (global and local) were used. Note that levels were revealed one at a time, thus allowing separation of their neural responses. This is not possible with traditional Navon figures. C) On the top the order in which the stimulus-blocks were shown. One example stimulus-block is expanded at the bottom, in which the global ‘E’s alternated with the mask. Participants detected occasional deviant shapes (in this example the last letter).

Previous work with these stimuli shows that it is easy to recognize consecutive stimuli of arbitrary shapes within one level (even at fast rates), but it is difficult to do so for both levels at once (Iglesias-Fuster et al., 2015). Furthermore, shifting attention between levels takes time, especially from global to local shapes in typical subjects (Valdés-Sosa et al., 2014). This makes sense, since it is improbable that each level of an object has its private neural mechanism for shape recognition (Tacchetti et al., 2018). Convergence of inputs from different levels on a common processor is more plausible, which would entail competition for common neural resource (Chelazzi et al., 1993). To reduce this competition, irrelevant levels must be filtered out. Fixing attention at the same level -when possible-would be advantageous. In fact, this type of attentional control can fail, producing illusory shape/level perceptual conjunctions (Hübner and Volberg 2005), and is atypical in conditions such as autism (White et al., 2009) and Alzheimer’s disease (Jacobson et al., 2005; Slavin et al., 2002). Thus cortical representations of the global/local levels, agnostic about shape, are needed to guide attention steadily to one of these tiers while ignoring the other (which makes H3 attractive).

Here, task-fMRI activity was measured in participants observing these modified Navon figures (Figure 2C). Multivariate pattern analysis (MVPA) was performed to search for cortical areas carrying shape-invariant level, or level-invariant-shape, information. We choosed a “searchlight” approach (Kriegeskorte et al., 2006), that allows an exhaustive whole brain exploration without prior assumptions on the location of the information. To detect invariance with MVPA, cross-classifications of each attribute (shape or level) must be used in order to see which activity patterns generalized across the other attribute (e.g. to see if the dissimilarity in fMRI patterns between global and local levels are equivalent across shapes) (Kaplan et al., 2015).

Each of the hypothesis about level information in higher-order cortex outlined above implies a different outcome in these MVPA. H1 entails a failure to reveal shape-invariant level information in any higher-order visual area, whereas H2 implies that level/shape information is present to the same degree in all of these areas. H3 -separate pathways- predicts that level and shape information -invariant to each other- are distributed unequally over the cortex. To anticipate our results, H3 was the most valid hypothesis in VTC as well as in other visual regions, in both hemispheres. Shape information was stronger in object-selective cortex. Interestingly, areas potentially carrying invariant information about level overlapped scene-selective cortex, although our stimuli were clearly objects/letters.

The relationship between the cortical patches carrying level/shape information was examined with functional connectivity (Friston, 2011), gauged by the correlation between their fMRI activity over time. We found stronger connectivity between areas with the same (compared to different) level/shape specialization. Due the special role of VTC in visual processing, a more detailed analysis was carried out in this region, testing different models of the topology of its connections. These tests suggested two independent caudal-rostral streams (each preferentially carrying either shape or level information), an outcome that also supported H3. Our results serve to better typify the networks involved in visual recognition, and suggest independent parallel neural pathways specialized for each attribute, especially in VTC.

## Materials and Methods

### Experiment

#### Participants

Twenty-six students from the University for Electronic Science and Technology of China (UESTC) participated in the main experiment (ages ranged from 18 to 28 years: mean = 22.5, std= 2.72; 9 females). All had normal (or corrected-to-normal) vision and were also right handed except for two cases. None had a history of neurological or psychiatric disease. The experimental procedures were previously approved by the UESTC ethics committee, carried out in accordance with the declaration of Helsinki, with participants giving written informed consent.

#### Stimuli and task

Stimuli were projected onto a screen in the back of the MRI scanner, viewed through an angled mirror fixed to the MRI head-coil, and generated using the Cogent Matlab toolbox (http://www.vislab.ucl.ac.uk/cogent.php). Modified Navon figures were built as follows. A matrix of white lines (about 2.0° wide and 5.3° high), on a black background, was used as a baseline stimulus. This matrix was built out of 15 small placeholder elements shaped like ‘8’s (spanning visual angles about 40′ wide and 1° 3′ high). Local letters were uncovered by eliminating selected lines within 10 out of 15 possible placeholders, whereas global letters were uncovered by completely eliminating several placeholders (see Fig. 2B). At both levels the letters ‘E’ and ‘U’ were presented. Small deviations in letter-shapes were used as infrequent oddball stimuli (see last letter in Fig 2C).

Each stimulus-block was initiated by a 1 sec cue (’Global’ or ‘Local’), that directed attention to the level at which the letter was to be unveiled. This was followed by the presentation of the baseline mask for 19 sec (see Fig. 2C and SI movie for examples). Then, the letter (at a fixed level) selected for each block was repeatedly presented 10 times, each instance lasting 1 sec and separated from the others by the reappearance of the baseline mask also for 1 sec. To encourage attention to the stimuli, participants were asked to count the number of oddball letters within a block (which occurred either 0, 1, or 2 times per block in equal proportions and a random places in the stimulus sequence). Note that neural responses to each block was triggered by the repeated switching between a letter and the mask, which would have weakened the contribution of elements common to the two. Blocks finished with a 4 sec ‘respond’ signal, with participants reporting the number of oddballs via a MRI-compatible button pad (detection accuracy in all participants were above 85%). Thus each block of recurring letters lasted 20 sec, and was separated from the next one by another 24 sec.

The experiment was divided into runs, each lasting 8.8 minutes, and separated by 1 to 2 minute breaks to allow the participants to rest. Each run contained 12 stimulus-blocks, that were 3 repetitions of the sequence of blocks containing the letters global ‘U’, global ‘E’, local ‘U, and local ‘E’ in that order. Most participants completed 5 runs, except two who only completed four. Consequently each type of stimulus-block was repeated a total of either 15 or 12 times in the experiment.

#### Data acquisition

Recordings for the experiment were obtained with a GE Discovery MR750 3T scanner (General Electric Medical Systems, Milwaukee, WI, USA) at UESTC, using an 8 channel receiver head coil. High-spatial resolution (1.875 × 1.875 × 2.9 mm) functional images (fMRI), with 35 slices covering all the head except the vertex (no gaps), were acquired. A T2*-weighted echo planar imaging sequence was used with the parameters: TR=2.5s; TE=40 ms; flip angle=90^○^ and acquisition matrix=128 × 128. There were 135 images per run. The initial 5 volumes were discarded for all runs, to stabilize T1 magnetization. A 262 slice anatomical T1-weighted image was also obtained with the following parameters: voxel size=1 × 1 × 0.5 mm; TR=8.10 ms; TE=3.16 ms; acquisition matrix=256 × 256; and flip angle=12.

#### Image preprocessing

Preprocessing was carried out with SPM8 (https://www.fil.ion.ucl.ac.uk/spm/). The functional scans were first submitted to artifact correction using the ArtRepair toolbox (http://cibsr.stanford.edu/tools/ArtRepair/ArtRepair.htm), thus repairing motion/signal intensity outliers and other artifacts (including interpolation using nearest neighbors for bad scans). Then slice-timing, head motion correction (including extraction of motion parameters), and unwarping procedures were applied.

White matter and pial surfaces were reconstructed from each T1-weighted image using Freesurfer software (http://surfer.nmr.mgh.harvard.edu), then registered to the FsAverage template, and subsequently subsampled to 81924 vertices (https://surfer.nmr.mgh.harvard.edu/fswiki/FsAverage). The mid-gray surface was calculated as the mean of white and pial surfaces vertex coordinates. Each T1 image was co-registered with the mean functional image, and the transformation matrix was used to project the mid-gray surface into each subject’s functional native space. Volume BOLD signals were interpolated at the coordinates of the mid-gray vertices, producing surface time-series (without spatial smoothing). Time series were high-pass filtered with cutoff of 0.008 Hz, their means and linear trends removed, and were also normalized (z-scored), all of which was performed for each run separately. A general linear model (GLM) was fitted to the time series of each surface vertex using a regressor for each stimulation block (i.e. square-waves convolved with the canonical hemodynamic function), in addition to the head movement parameters and the global mean of the white matter signal as nuisance covariates. The beta parameters estimated for each block (trial) were used as features for MVPA, whereas the residual time series or background activity (Al-aidroos et al., 2012) were used to perform the functional connectivity analysis.

### Searchlight MVPA for decoding invariant information for level and shape

The overall design of the analyses, of fMRI data and their relation to the questions addressed in the introduction are shown in Figure 1B. Multivariate pattern analysis has the advantage to be more sensitive than traditional univariate activation, due to its ability to find groups of nodes with weak activation, but consistent across experimental conditions. Here, the presence of invariant level/shape information in the fMRI was gauged through a searchlight approach (Kriegeskorte et al., 2006). The searchlights are multiple, overlapping, small regions of interest, which exhaustively cover the cortical surface. A classifier is training and tested in each searchlight for each subject and then submitted to group analysis.

Cross–classification inside MVPA was employed to ascertain invariant level and shape information in each searchlight (Kaplan et al., 2015). Shape-invariant level information was considered to be present if letter level was predicted accurately (i.e. above chance) for one letter by a classifier trained on exemplars of the other letter (Fig. 3A). This would imply patterns for level indifferent to letter shape. Conversely, level-invariant shape information was considered to be present if letter identity at one level could be predicted accurately by a classifier trained on exemplars from the other level. This would imply multivariate patterns for shape tolerant to large changes in physical format between different levels (Fig. 3B).

**Figure 3.**
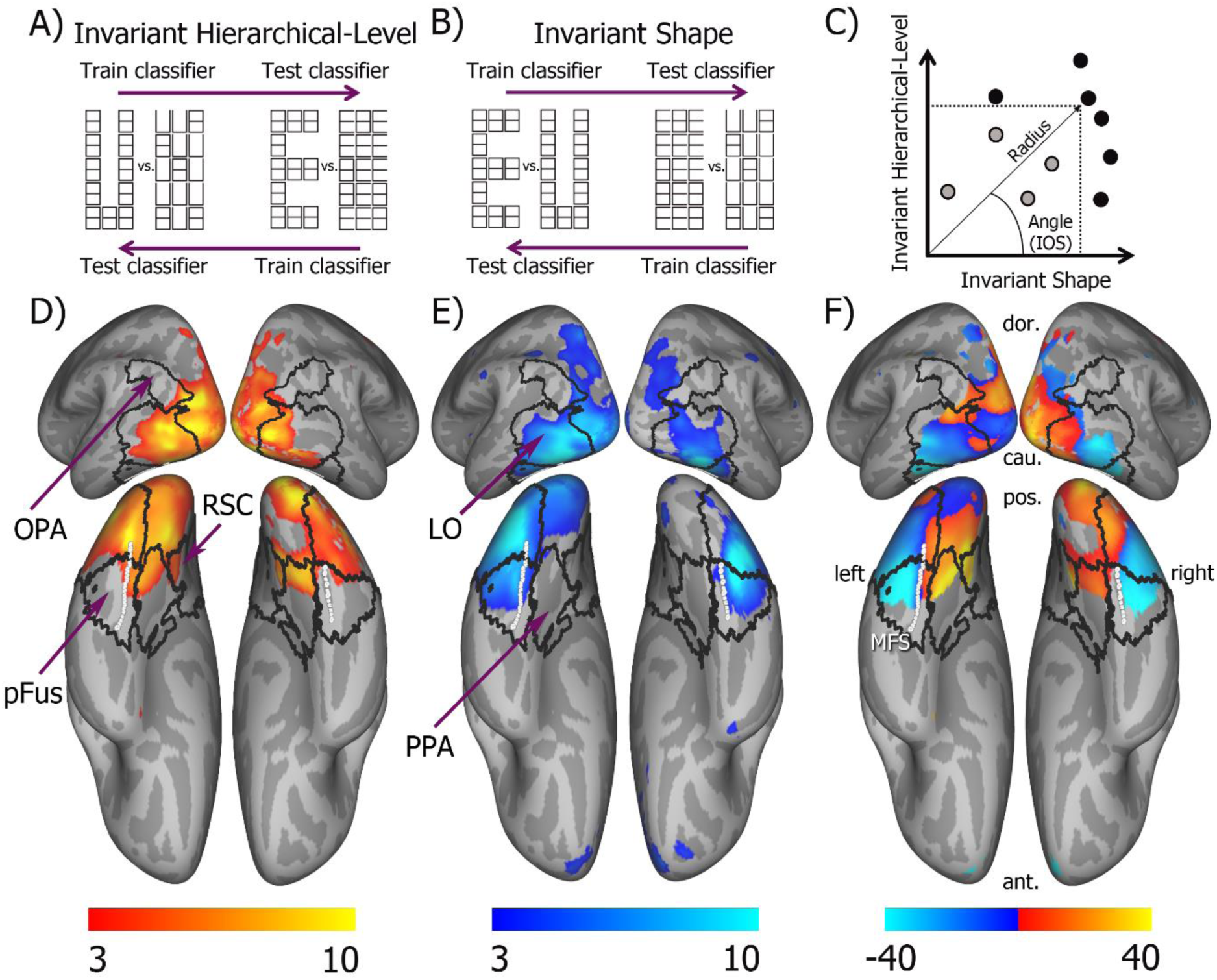
Searchlight maps. A) Design of cross-classification procedure for shape-invariant level. B) Design of cross-classification procedure for level-invariant shape. For A and B accuracies of classification in the two directions were averaged. C) IOS defined as the polar angle between the accuracies of level and shape classification at each searchlight (minus 45 degrees), which we illustrate with an arbitrary number of nodes. Dotted lines are FDR thresholds for each cross-classification. D) Searchlight map for shape-invariant hierarchical-level decoding, showing group Z-scores of above-chance classification (thresholded FDR q=0.05). E) Corresponding searchlight map for level-invariant shape decoding. F) Group IOS map. Only searchlights informative for at least one cross-classification are shown (drawn in C as black circles in this toy example). Also, borders of scene- and object-selective areas are shown in black lines. Object-selective areas: LO lateral occipital; pFus posterior fusiform. Scene-selective areas: PPA Para-hippocampal place area; RSC retrosplenial complex; OPA occipital place area. The MFS is depicted in white dotted lines. Acronyms: dor.: dorsal; cau.: caudal; pos.: posterior; ant.: anterior

Also, we introduced an Index of Specialization (IOS) as a measure of the relative presence of level -or shape- invariant information in each searchlight. IOS was expected to reveal spatially segregated cortical sectors according to H3 but not H1 or H2.

#### Searchlight analysis

Discs with 10 mm radii of geodesic distance were defined around all 81924 vertices (mean number of nodes: 186; range: 53-471) by means of the fast_marching toolbox (https://github.com/gpeyre/matlab-toolboxes). Then, MVPA was performed in each subject on the beta parameters for the set of vertices within each searchlight using a fast Gaussian Naïve Bayes classifier developed for searchlight computation (Ontivero-Ortega et al., 2017) in a leave-one run out cross-validation scheme (for 5 runs, 12 trials was used for training and 3 trials was used for testing, per condition in each iteration of cross-validation). In turn, for cross–classification four different classifications were employed, two for level and two for shape: Level (train E global and e local, test U global and u local; train U global and u local, test E global and e local), Shape (train E global and U global, test e local and u local; train e local and u local, test E global and U global).

For group analysis, individual accuracies in each searchlight were averaged across folds and in both directions of the cross-classifications (for level and for shape), resulting in two main searchlight maps for each subject. The resulting averages were then logit transformed (logit-acc), and assigned to the central node of each searchlight. To generate the final searchlight maps, these two logit-acc datasets (for level and for shape) were independently submitted to a group t-test vs chance level (accuracy=50%, logit-acc=0). A false discovery rate (FDR) threshold of q=0.05 was used to control the effects of multiple comparisons in both maps, based on estimation of empirical Gaussian null distributions (Schwartzman et al., 2009). Note that the use of cross-classification mitigates limitations of group-level t tests for MVPA (Allefeld et al., 2016) due to the positively skewed accuracy distributions which for binary classifications should generally larger than 50%. However, in the case of cross-decoding accuracies below 50% can be expected if there are interactions between factors (i.e.: when patterns are systematically different across a secondary factor).

However, searchlights may be informative (i.e. allow decoding) either because the local patterns of activity differ across conditions, or because response amplitudes differ (without pattern change), or perhaps because of both (Coutanche, 2013). A traditional univariate analysis should be enough for revealing the contribution from response amplitudes. Yet, to test if local patterns contribute to the MVPA results, for each trial the mean of the betas in a searchlight was removed (i.e. the data was centered) before MVPA. To test if the amplitudes contribute to the MVPA results, the beta values in each searchlight were replaced by their average across nodes before MVPA (i.e. the data was smoothed, similar to univariate analysis). This corresponds to spatial smoothing with a cylindrical (instead of a Gaussian) window. These transformed maps were analyzed as described above. The correspondence between different types of maps was assessed with the Dice coefficient (Dice, 1945).

#### Index of Specialization (IOS)

The degree of specialization for level/shape invariant information was characterized in each searchlight by an index of specialization (IOS, Fig. 3C), which reflected the relative accuracy of the two types of cross-decoding. After expressing logit-acc for the two attributes in polar coordinates (negative values were substituted by zero), IOS was defined as the polar angle minus 45 degrees. Therefore −45 degrees corresponded to maximum specialization for shape information, 45 degrees to maximum specialization for level information, and zero to no preference. IOS was only calculated in informative searchlights (i.e. in which at least one of the t-tests vs chance was significant). This measure is similar to the activity profiles of different stimuli (expressed on spherical coordinates) that have been used in clustering voxels (Lashkari et al., 2010), although here informational-instead of activity-profiles were employed.

Henceforth, two non-overlapping functional regions (ROI) were defined in each hemisphere: a shape-invariant level ROIs considering all nodes with IOS>10, and a level-invariant shape ROsI with IOS<-10. This arbitrary value was selected to rule out values close to zero (i.e. with no clear preference for level or shape processing). To help describe the searchlight maps, nodes corresponding to the mid-fusiform sulcus (MFS) (Weiner, 2018) were identified manually on the FsAverage surface. Also, the VTC region was defined as all the vertices included in fusiform gyrus, the lingual gyrus, and the lateral occipito-temporal, collateral, and transverse collateral sulci, according to the Freesurfer atlas (Destrieux et al., 2010), but only posterior to the rostral tip of MFS.

### Comparison of searchlight IOS map with category-selective functional localizer maps

Regions where level and shape were potentially invariant to each other (selected from the IOS map) were compared to well-characterized functional regions of interest (fROIs) using population templates of visual category-specific cortex. This allowed describing our results in the context of previously published work.

Scene-selective areas (Zhen et al., 2017), defined by the contrast of scenes vs visual objects, included the para-hippocampal place area (PPA), the retrosplenial complex (RSC) and the occipital place area (OPS). Object selective areas, defined by the contrast of objects vs scrambled objects, (http://www.brainactivityatlas.org/atlas/atlas-download/), included two portions of the lateral occipital complex (LOC): the lateral occipital (LO) and the posterior fusiform areas (pFus). Given the possibility that level could be represented by attentional-zoom on retinotopic presentations, an atlas of 25 visuotopic maps (Wang et al., 2015) was also used. Since we could not calculate our IOS index on the data from these probabilistic maps, we additionally carried out a face/house fMRI localizer experiment in an independent group of participants at the Cuban Centre for Neuroscience (see SI). An analog of the IOS was calculated on the data from this control experiment.

### Background activity connectivity analysis

This analysis aimed to characterize the functional connectivity (FC) of the IOS level/shape invariant regions. We used the background activity (BA) (Al-aidroos et al., 2012) time series to do this. This estimation strips-out the contribution of the stimulus-locked response (stimulus-driven connectivity), which is time-locked to stimulus perceptual availability, and looks at between areas state-dependent connectivity. FC was estimated with the Pearson correlation coefficient.

#### Comparison of the FC between regions with similar and dissimilar IOS

We first tested if the FC was stronger between areas with the same specialization, rather than dissimilar specialization. The regions for invariant level and shape (excluding V1 and V2) were each divided into spatially separated clusters (contiguous surface nodes were identified using the clustermeshmap function from the Neuroelf toolbox (http://neuroelf.net/)). Only clusters containing more than 100 nodes were considered. In each subject, BA time series were averaged across nodes in each cluster. Partial correlations were calculated between all pairs of these IOS-defined clusters (controlling for the correlation with the other clusters). Cells of the resulting partial correlation matrices were averaged within each individual after grouping according to two main effects: Pair-similarity (shape-shape, level-level or shape-level) and Hemisphere (clusters in the same vs. different hemisphere). A repeated measure ANOVA was then performed on the reduced matrix values after a Fisher transformation, using these main effects.

#### Prediction of IOS parcellation from FC in VTC cortex

To verify the relationship between FC and cortical specialization we carried out a comparison between a functional connectivity parcellation and IOS maps. We limited this analysis to the VTC region for each hemisphere separately, where side-by-side two different functional domains were defined by the mid fusiform sulcus division. First, a vertex-wise FC matrix was estimated in each individual by calculating the Fisher transformed Pearson correlation between all cortical nodes in VTC. The group-level matrix was thresholded with a t-test against zero that was corrected for multiple comparisons (p<0.05, Bonferroni corrected). This mean FC matrix was then partitioned into two new FC-defined clusters using spectral clustering method (Von Luxburg, 2007). The hypothesis was that two regions specialized (for shape and level) would emerge from the cluster method. Two ensure stability, the clustering process was repeated 100 times and a consensus matrix C (Lancichinetti and Fortunato, 2012) was built based on the different parcellations across the iterations. The final two FC-defined clusters were generated over C using again the clustering method, and the dice coefficient between these clusters and the IOS regions was calculated for both hemispheres, as well as the mean IOS across their nodes.

#### Tests of models of FC structure in VTC

Finally, we studied the topology of FC inside VTC, testing several theoretical models that could explain partial correlations within this region, with an approach analogous to that used in Representational Similarity Analysis (Kriegeskorte et al., 2008). In each hemisphere, the VTC ROI was restricted by excluding non-informative-searchlights, as well as the V1 and the V2 regions. The BA time-series of each node was spatially smoothed on the surface by replacing it with the average of all nodes in the (10 mm radius) searchlight surrounding it. Each restricted VTC was divided in 10 patches along the caudal-rostral direction by a k-means partition of the Y coordinates of the surface nodes. The two FC-defined clusters in each hemisphere described above were then subdivided with these 10 patches. This yielded in both hemispheres 8 and 9 patches for each cluster respectively (Fig. 5C). A patch-wise observed FC matrix was estimated in each participant, by calculating partial correlations (to remove spurious or indirect associations) with the averages of BA within patches.

Alternative theoretical models (expressed as patch-wise matrices) were built as explanations of the pattern of connectivity in VTC. These models were: a) connectivity by simple spatial proximity (indexed by the matrix of average between-node geodesic distances over the cortex for all pairs of patches); b) original FC-defined cluster membership; c) patch contiguity in the caudal-rostral direction (CR adjacency); and d) lateral patch contiguity (lateral adjacency). The models and the patch-wise FC matrices were vectorized. Multiple regression was carried out separately for the two hemispheres in each subject using the theoretical correlation values (corresponding to each model) as the independent variables and the observed connectivity values as the dependent variable (*obsCorr = Intercept + B_1_*geodesic distance + B_2_*cluster + B_3_*CR-adjacency + B_4_*Lateral-adjacency*). The intercept was considered a nuisance variable. The resulting beta values (*B_i_*) across participants were submitted to a random effect t-test against zero. The effect size of each predictor was estimated by bootstrapping the test (*n=1000*). This allowed selection of the theoretical matrix most similar to the observed patch-wise FC matrices.

## Results

### Information about level and shape are carried by different cortical regions

Invariant level and shape searchlight maps (Fig. 3D, E and Supplementary Fig. S2A, B) overlapped only moderately (Dice coefficient = 0.5). Referenced to anatomical landmarks (Destrieux et al., 2010), level information was most accurately decoded in the occipital pole, but also in the medial portion of the fusiform gyri and the lingual gyrus, the collateral and traverse collateral gyri, as well as medial occipital areas. Conversely, shape information was concentrated in the lateral occipital and posterior lateral fusiform gyri, as well as the lateral-occipital sulci.

The centered searchlight maps (Fig. 4A, B and Fig. S3A, B), which show decoding based only on patterns, were very similar to the original (untransformed) maps (Dice coefficients: for level=0.82; for shape=0.74). Thus local patterns contribute to information for both level and shape at most sites. Smoothed maps, which show decoding based only on amplitude (Fig. 4D, E; and Fig. S4A, B) were similar to the original maps for shape (Dice coefficient=0.69), but were different fin the case of level (Dice coefficients for level=0.17). This discrepancy is due to the fact that the smoothed map for level was only informative in caudal, but not rostral (higher-order) visual areas. Thus, information about level was carried by amplitude exclusively in caudal early visual cortex.

**Figure 4.**
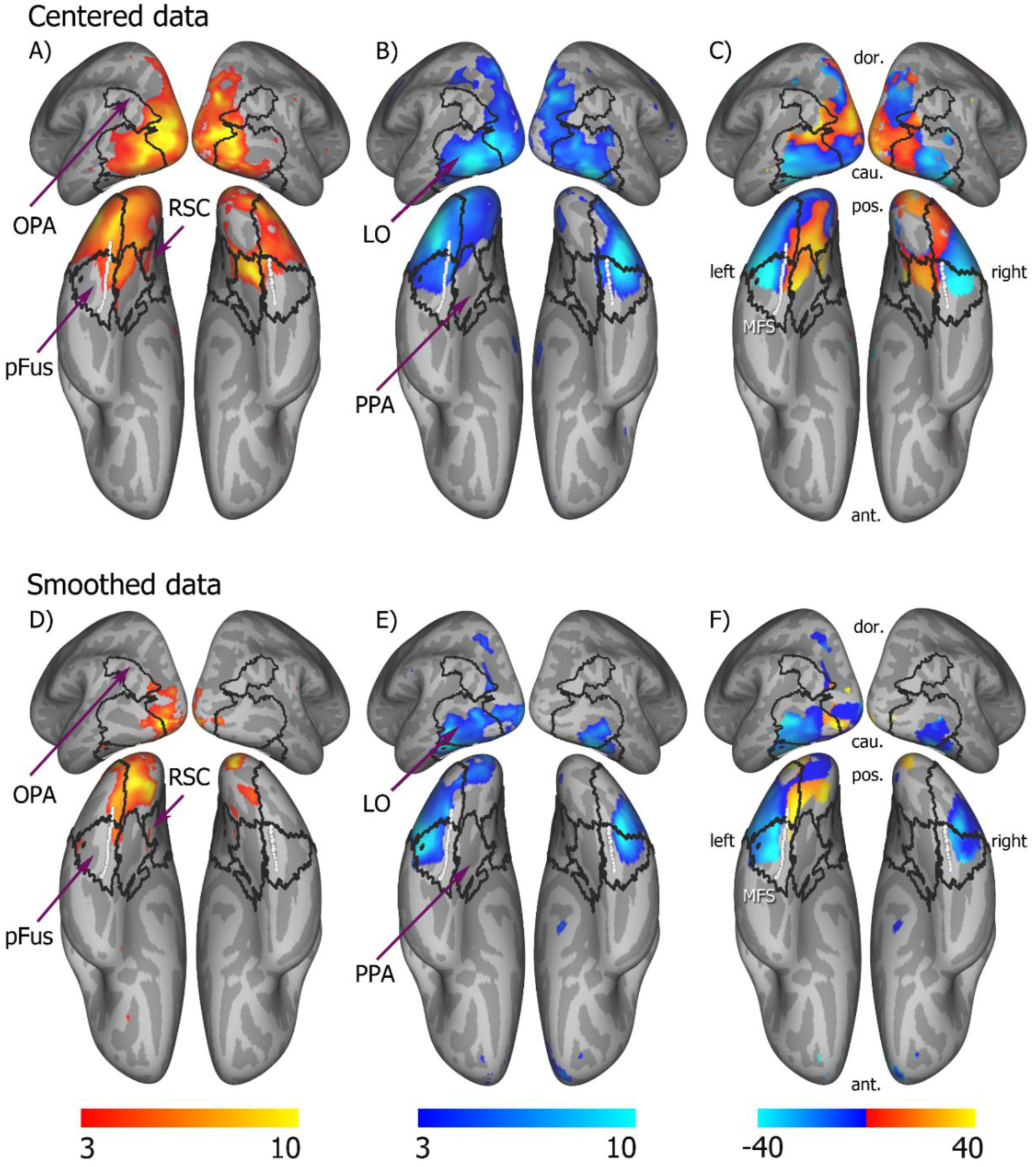
Searchlight maps for centered and smoothed data. Centered maps: A) Shape-invariant level map; B) Level invariant shape map; C) IOS map. Smoothed maps: D) Shape-invariant level map; E) Level invariant shape map; F) IOS map. ROI names and orientation conventions as in Fig. 2.

The IOS map (Fig. 3F and Fig. S2C) confirmed the spatial segregation level or shape information preference. Singularly, a sharp boundary at the mid-fusiform sulcus (MFS), divided VTC into areas with different informational profiles: a lateral area more specialized for shape and a medial one more specialized for level. The fact that level and shape were carried by different (albeit overlapping) cortical areas rebutted H1-2 and vindicated H3. The centered IOS (Fig. 4C and Fig. S3C) and original IOS maps were highly similar (Dice coefficient = 0.83), however the smoothed IOS (Fig. 3F and Fig. S4C) and original IOS maps overlapped less (Dice coefficient =0.48).

### Location of IOS regions relative to ROIs from probabilistic atlases

Searchlights preferring shape-invariant level information occupied (see Supplementary Table S1) the posterior portion of the PPA in both hemispheres, with a slight overlap with the ventral-posterior portion of the RSC and the OPA. Additionally, overlap was found with the superior posterior portions of LO division in both hemispheres. Referred to visuotopic maps (see Supplementary Figure S5 and Table S1), shape-invariant level information was found in PHC1, VO2, V3 and ventral V3 as well as V3A/B and superior posterior parts of LO1-2 in both hemispheres. PHC2, ventral V2, and V1 on the right, as well as VO1 on the left side were also involved. Note that PHC1-2 corresponds to the retinotopically-organized posterior part of PPA. Similar results have been obtained in previous mapping of scene-selective ROIs onto retinotopic areas (Epstein and Baker, 2019; Malcolm et al., 2016; Silson et al., 2016). Importantly, whereas mean-corrected and smoothed decoding of shape-invariant level were both accurate in all early visual (V1-V3, hV4), only mean-centered decoding was found in VO1-2 and PHC1-2. This indicated that the visual features underlying level information shifted in the caudal to rostral direction.

Conversely to level, and unsurprisingly, searchlights preferring level-invariant shape information in both hemispheres overlapped the LOC, specifically antero-ventral LO and most of pFus, as well retinotopic areas LO1-2 and TO1-2 (see Supplementary Figure S5 and Table S1). These areas are well known to be selective for visual objects (Kourtzi and Kanwisher, 2000). Note that the border between level and shape dominance corresponded better to the MFS than to the borders of the PPA, LO, and pFus fROIs from the atlas. This motivated an application of our IOS measure to data from a face/house localizer experiment (described in supplementary information). In addition to verifying an overlap of the level IOS regions with scene (i.e. house) selective cortex, as well as a more moderate overlap of shape IOS regions with face-selective cortex, the IOS map for this experiment showed a sharp functional divide exactly at MFS (see Figure S6). In other words, when IOS was used to characterize the border between scene -and object-selective cortex, we observed the same boundary at MFS as found for level/shape.

### Areas specialized for level and for shape belong to independent pathways

Pairs of IOS-defined clusters with similar level/shape specialization (IOS) had stronger connectivity than pairs with different specialization. The repeated measure ANOVA showed a highly significant effect of Pair-similarity (F(2,50)=169, p<10^−5^) on the partial correlation (Fig. 5A), which was driven by lower values for shape-level than for shape-shape and level-level (F(1,25)=285, p<10^−5^) in both hemispheres. No effect of Hemisphere was found, although it interacted significant with Pair-similarity (F(2,50)=7, p<0.002). This interaction corresponds to significantly larger partial correlations (F(1,25)=23, p<10^−3^) for level-level than for shape-shape pairs of different hemisphere.

**Figure 5.**
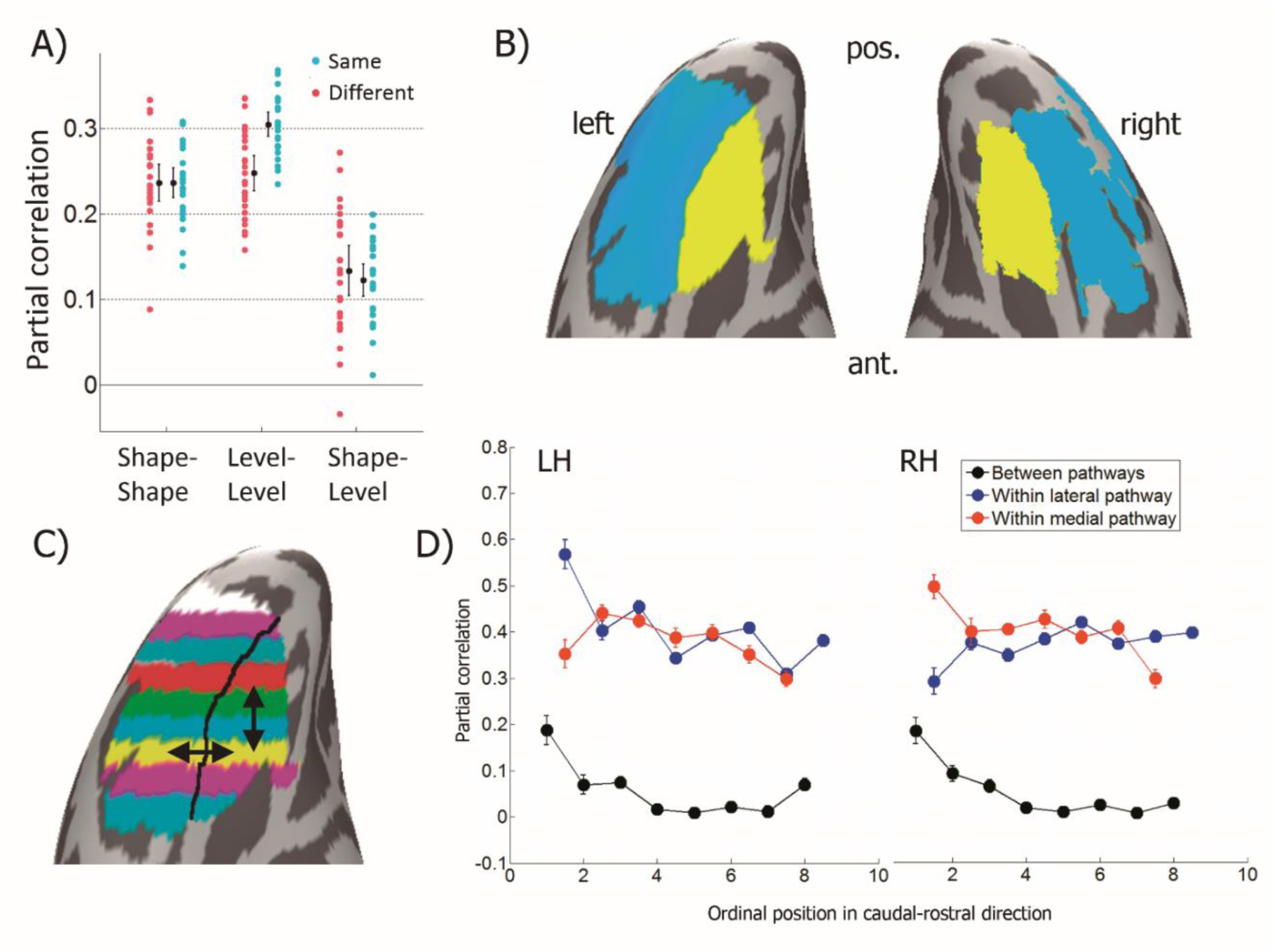
Functional connectivity analysis. A) Scatterplots of mean partial correlations between initial clusters as a function of their similarity in level/shape specialization and if they are in the same or different hemispheres. B) The two clusters obtained by Spectral Clustering method for each hemisphere. C) Subdivision of the latter clusters into parallel bands (here and in D-E shown only in the left hemisphere). Arrows show examples of bands that are adjacent in the caudo-rostral direction and the lateral direction. D) Partial correlations between patches in VTC for each hemisphere; red (lateral cluster) and blue (medial cluster) plots correspond to values between consecutive bands in the anterior-posterior direction, whereas black plot corresponds to values between adjacent patches in the two clusters. In A and D, dots and whiskers respectively represent means and standard errors of means.

The ANOVA showed that IOS similarity predicted FC, so we asked if FC could predict the IOS parcellation. In each hemisphere the FC-based parcellation divided VTC into two compact areas that differed in mean IOS values: a lateral region (mean IOS = −13.7), and a medial region (mean IOS = 20.6) (Fig. 5B). The clusters derived from FC analysis corresponded very well to the segmentation produced by the IOS map. The medial cluster overlapped level-informative areas, whereas lateral cluster overlapped shape-informative areas. In other words, FC predicted cortical specialization. Fig. 5D shows the plots of the group-mean partial correlations between adjacent patches in the caudo-rostral direction, and between adjacent patches in the lateral direction. It is clear that the patches in the caudo-rostral direction were more connected (i.e. had higher partial correlations) than those in the lateral direction (which crossed the MFS).

### Topology of connections in VTC is related to informational specialization

The topology of functional connectivity (FC) patches in VTC was tested more formally by comparing the amount of variance of the observed partial correlation matrix associated with each of the four theoretical matrices outlined above (Fig. 6A). The observed correlation matrix (Fig. 6B) was best explained by the model specifying adjacency in the caudo-rostral direction in both hemispheres, consistent with the effects described above. The effect size of this organizational model was much larger than for the other three models (Table I). This suggests two parallel functional pathways within VTC, each coursing in a caudal-rostral orientation.

**Figure 6.**
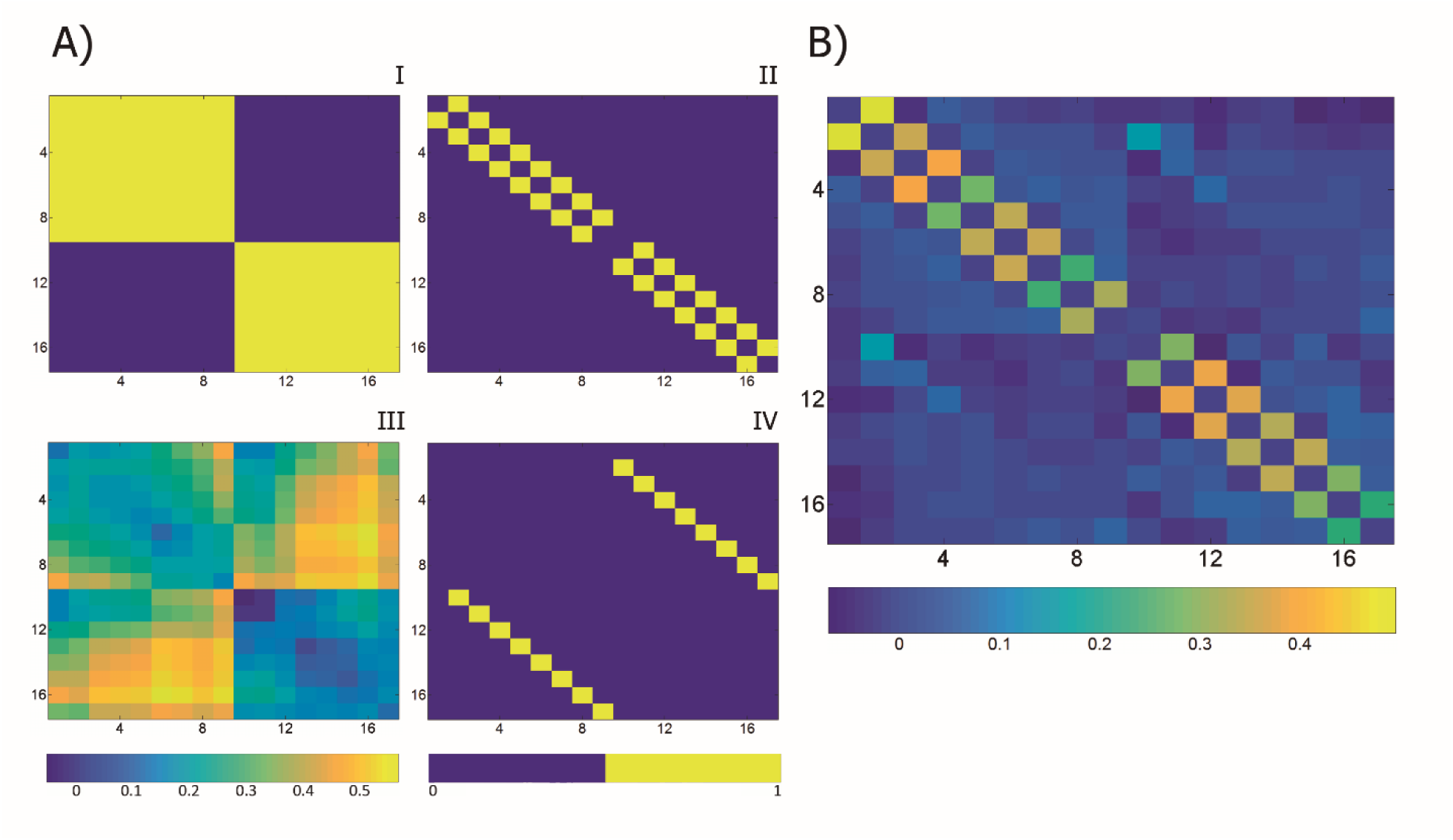
Topology of FC in VTC. A) Competing models: I - Cluster membership; II - Caudo-rostral within-cluster adjacency between bands; III - Geodesic distance between all pairs of bands; and IV - Lateral adjacency between bands from different clusters. B) Experimentally observed group-mean partial correlation matrix.

**Table I.**
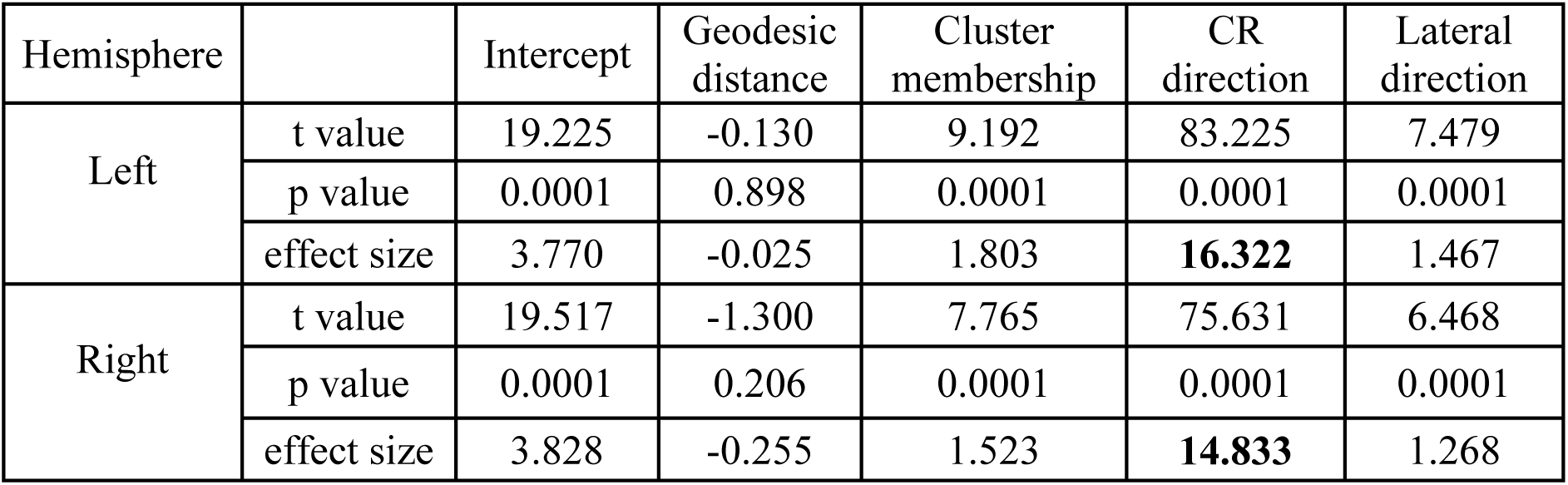
Assessment of competing models as predictors of between-band FC in VTC (multiple regression analysis).

In summary, similarity in the cortical specialization of cortical areas for level vs. shape (indexed by the IOS) predicted their FC. On the other hand, FC parcellated the VTC into areas that were specialized for either level or for shape. These results buttress H3, which posits that hierarchical level and shape of visual objects are processed in separate neural circuitry, and negates H1 and H2.

## Discussion

We presented three alternative hypothesis in the introduction. Two findings validated H3, which posits distinct pathways within VTC, each better specialized in processing either shape or hierarchical-level. The first finding was that cross-decoding of level and of shape (evincing invariance to each other) was unevenly distributed over the cortical surface. Shape-invariant level decoding was better in areas previously characterized as scene-selective. Oppositely, level-invariant shape decoding was better in regions reported to be object- and face-selective. The boundary between regions with different informational profiles was marked sharply by the MFS, thus extending the list of functional and anatomical parcellations delimited by this sulcus (Grill-Spector and Weiner, 2014). Second, functional connectivity analysis indicated that cortical patches having a similar informational specialization were more strongly interconnected than those that had divergent specialization, with a pattern within VTC suggesting two parallel pathways, each placed on a different side of the MFS.

The demonstration of level information in scene-selective cortex was possible due to combination of novel stimuli with MVPA. To our knowledge, MVPA has not been used in previous fMRI studies of traditional Navon figures (Han et al., 2002; Weissman and Woldorff, 2005), but even if it were applied, shape-invariant level and level-invariant shape information would go undetected with these figures. With traditional stimuli, global and local shapes are presented at once, therefore their activity patterns cannot be separated. This is unfortunate because the cross-decoding approach used to diagnose invariance to secondary attributes depends on this separation. In our modified Navon figures the two levels are presented separately over time, it is therefore possible to estimate the activity patterns separately for global and local target shapes, overcoming this obstacle.

Although the overall shape of our local letters (which occupied a full rectangle) was different than that of our global letters (in which the rectangle had gaps), this distinction in overall configuration probably did not contribute to the test for level information. The fMRI activity patterns that mattered for the MVPA were triggered by the discrepant features in the rapid alternation between the letters and the background mask (see Figure 2C). The rectangular configuration did not change in the back and forward switch between local letters and mask. The effective stimulation in this case consisted in circumscribed offsets of lines, which were unevenly distributed within the rectangular area. Furthermore the precise location of these local changes were different for the two variants of local letters, as were the location and orientations of the gaps in the two variants of global letters. Hence what we dubbed cross-decoding of level could not have been based on the distinction between the overall geometric outline of the stimuli.

Decoding of hierarchical-level from scene-selective cortex seems counter-intuitive. Navon figures are visual objects not places. However, features implicated in scene processing (Groen et al., 2017) could also be used to code level. The low-level visual feature of spatial frequency serves to distinguish scenes (Andrews et al., 2015; Berman et al., 2017), and is also important in selective attention to hierarchical levels (Flevaris et al., 2014; Flevaris and Robertson, 2011; Valdés-Sosa et al., 2014). In mid-level vision, clutter (Park et al., 2015), amount of rectilinear contours (Nasr et al., 2014), and statistics of contour junctions help distinguish between scenes (Choo and Walther, 2016). These features also differ markedly between the global/local levels of Navon figures (e.g. see Figure 1 in (Flevaris et al., 2014)). Topological properties, such as number of holes was different in the global/local stimuli, which is perhaps related to clutter and statistical of contour junctions. Finally high-level features such as spatial layout (in conjunction with object content) can be used to guide the navigation of gaze and attention within a scene (Malcolm et al., 2016). It is possible that the same features would allow attention to be navigated within Navon figures. The overall position, size, and orientation of gaps in our figures are feasibly related to properties of spatial layout such as such as openness and expansion that have been examined in research on scene selectivity (Oliva and Torralba, 2001).

Finding shape information that is invariant to other attributes in object-selective cortex was otherwise unsurprising. Previous MVPA studies have shown that the shape of visual objects can be reliably decoded in LOC with tolerance to variations in their position (Cichy et al., 2011; Stokes et al., 2011; Williams et al., 2008), size and viewing angle (Eger et al., 2008a, 2008b), and rotation (Christophel et al., 2017). Also if the overall shape is maintained, fMRI-adaptation for the same object survives in LOC despite interruptions of its contours (Kourtzi and Kanwisher, 2001). It is not clear how invariance of shape to global/local level is related to invariance for these other attributes.

The most evident explanation for shape-invariant level decoding – as mentioned in the introduction- could simply reflect the use of different attentional-zooms in the visual field for global and local shapes. Global shapes are larger, and have lower spatial frequency content than local shapes, which corresponds with the tuning of more peripheral sectors of visuotopic maps. Local shapes are smaller and have higher spatial frequencies than global shapes which corresponds with the tuning of foveal sectors (Henriksson et al., 2008). Congruent with this notion, two studies using mass univariate fMRI activation tests have shown that foveal sectors of early visual areas (V2, V3, V3A, and hV4) are more activated by attention to local -compared to global-stimuli (Rijpkema et al., 2007; Sasaki et al., 2001). The effect is not evident in higher-order visual areas.

Here, the smoothed searchlight maps are most similar to mass univariate fMRI activation tests. In these maps, consistent with the previous studies, shape-invariant level was found only at the foveal confluence of early visual areas (V1-3), but was absent in higher-order visual areas (VO1-2, PHC1-2). In contrast, in the latter areas, level information invariant to shape was found with the centered searchlight maps (which are pattern- but not amplitude-dependent). This suggests a change in code format between early visual areas and higher-order areas, with attentional-zoom only in the former group. However, in the latter areas the link between visuotopic mapping and anatomy is less direct and strongly modified by attention (Kay et al., 2015), making it more difficult to detect. Therefore, a rigorous test of possible change in level code format requires testing cross-decoding and pRF maps in higher-order areas within the same experiment.

There is another way attentional-zoom could underlie cross-decoding of level. The medial and lateral VTC have different retinal eccentricity biases, preferentially linked to peripheral/foveal stimuli and sectors of V1 (Baldassano et al., 2016b; Hasson et al., 2002) respectively. Switching attention between the global/local levels could potentially shift activity between medial and lateral VTC, thus allowing cross-decoding of level. Nevertheless, this scheme cannot explain why we found invariant-level decoding within searchlights restricted to only one sub-region (nor why decoding is more accurate in medial VTC).

Non-retinotopic coding of level is implied by two experimental findings from the literature. Attention to one level of a Navon figure selectively primes subsequent report of one of two superimposed sinusoidal gratings (Flevaris and Robertson, 2011). Attention to the global/local level respectively favors the grating with lower/higher spatial frequency, in apparent support of retinotopic coding. However, the relative (not the absolute) frequency of the two gratings determines this outcome (Flevaris and Robertson, 2016), which precludes coding by mapping onto retinal eccentricity. Moreover, although larger activations for big-compared to small-real world size objects have been found in medial-VTC, this effect is tolerant to changes in size on the retina. Both these phenomena suggest that a mechanism for extracting relative scale exists in the medial VTC, which could be used to represent shape-invariant level.

Our results suggest that the internal topology of VTC is dominated by rostro-caudal connections, with a division that maps onto level/shape specialization. A recent analysis (Haak and Beckmann, 2016) using functional connectivity patterns of twenty-two visuotopic areas (based on the same atlas used here, Wang et al., 2015) found a tripartite organization that is consistent to some extent with our results. One pathway lead from V1 into VO1-2 and PH1-2, largely overlapping our medial route. Another pathway lead into LO1-2 and TO1-2, partially overlapping our lateral route. Unfortunately in this study pFus connectivity (which looms large in our results) was not examined. Our results indicate that these pathways are organized to connect areas with similar informational specialization. This is congruent with other studies that show (Hutchison et al., 2014; Stevens et al., 2015) stronger FC between areas with the same, as compared to different, category-selectivity (e.g. faces, scenes, tool-use). Pathway segregation according to content (e.g. level vs. shape) allows implementation of divergent computations for different tasks.

Two studies with diffusion-tensor-imaging (DTI) tractography suggest an anatomical basis for the two functional VTC pathways posited here. One study (Gomez et al., 2015) found fiber tracts coursing parallel to each other in a caudal-rostral orientation within VTC. One of the tracts was found within face-selective, and the other within scene-selective areas (corresponding to our level/shape areas). In the other study, probabilistic tractography was used to cluster fusiform gyrus voxels based on their anatomical connectivity with the rest of the brain. The fusiform gyrus was divided into medial, lateral, and anterior regions (Zhang et al., 2016), the first two separated by MFS (also corresponding to our level/shape division).

Only the posterior portions of PPA, OPA and RSC (roughly corresponding to PHC1-2, VO1-2 and V3AB) potentially contain shape-invariant level information, contrariwise to their anterior sections. This limited overlap thus involved a posterior scene-selective sub-network found with FC (Baldassano et al., 2016a; Çukur et al., 2016; Silson et al., 2016). This sub-network, centered on the posterior PPA and OPA and not coupled to the hippocampus, would processes visual features needed to represent spatial layout. An anterior subnetwork containing the anterior portions of RSC and PPA (and strongly connected to the hippocampus), would play a role nonvisual tasks like memory for scenes, which would be poorly mobilized by our task thus explaining why level information is not detectable there. Furthermore, the posterior subnetwork would be biased toward static stimuli (our case), whereas the anterior network would be biased toward dynamic stimuli (Çukur et al., 2016).

Further work should take into consideration the following limitations of our study. We only included two shapes in the design, as in previous studies of shape decoding (Stokes et al., 2011), to improve the signal-to noise-ratio for estimation of multivariate patterns. The generality of our findings must be tested with a larger range of shapes and stimuli of different retinal sizes (to help answer determine if absolute or relative size contributes to the level decoding). With more diverse stimuli, representational similarity analysis (Kriegeskorte, 2008) could be used to better characterize level/shape representation in higher-order visual areas. Visual field mapping of pRFs (Dumoulin and Wandell, 2008) and level/shape cross-decoding must be performed in the same experiment to adequately test the attention-zoom explanation. Other limitations were that eye-movements were not controlled, and the face-house/contrast localizer was obtained from different subjects than the main experiment. Additionally, other patterns classifiers should be tested, since comparing their accuracy could give clues about differences in coding between areas (as found here by comparing smoothed and centered searchlight maps). Finally, although coding of level plausibly contributes to the representation of visual whole and parts, other sensory attributes probably involved should be examined.

Separate pathways for invariant level and shape have many functional implications. Shapes from different levels of the same object could compete for the same neuronal receptive fields (Chelazzi et al., 1993). This is illustrated by a study presenting two fast streams of our modified Navon figures, one global and the other local, at the same visual location (Iglesias-Fuster et al., 2015). Shape identification was more accurate when attention was focused at only one level than when it was split between two. To guide filtering of the irrelevant level, invariant information about level is required in higher-order visual areas. We posit that scene-selective cortex can play this role. Moreover, brief presentation of Navon figure leads to illusory shape/level conjunctions (Hübner and Kruse 2011; Hübner and Volberg 2005; Flevaris, Bentin, and Robertson 2010). Letters from the non-target level are fallaciously seen at the target level This implies that shape/level codes must separate somewhere in the brain (Hübner and Volberg, 2005), perhaps in the two VTC pathways described here. Thus independent coding of level and shape may facilitate sustained attention to a single level, but at the price of risking illusory conjunctions with impoverished attention.

### Conclusions

Our results refuted H1 since level information does not disappear from higher-order visual areas. H2 was largely rebutted because there was a significant divergence in the cortical areas preferring level and shape information. But consistent with previous work (Cichy et al., 2011; Golomb and Kanwisher, 2012; Hong et al., 2016; Rauschecker et al., 2012; Zopf et al., 2018), information about the non-preferred attribute does not disappear in the shape- and level-preferring pathways. However, a relative reduction of information about the non-preferred attribute was found in each pathway, which means a modified version of H3. This suggests computational circuitry specialized for achieving invariance for a primary attribute, geared to specific real-world demands (Peelen and Downing, 2017), that however does not completely discard information about secondary attributes.

The fact that shape-invariant level was potentially decodable from scene-selective cortex was an interesting result. Although Navon figures are clearly visual objects, they are not only that. If we accept the definition (Epstein and Baker, 2019) that we act ***upon*** objects, but act ***within*** scenes, it is possible to understand the dual nature of Navon figures. Shapes are acted ***on*** to recognize them, but are selected ***within*** the framework of their organization into levels. We postulate that coordinated use of invariant level and shape codes not only helps navigate attention between levels in Navon figures, but possibly within all real-life visual objects possessing wholes and parts.

## Acknowledgments

We thank the University of Electronic Science and Technology of China (UESTC) for funding and providing special equipment and facilities for this project. Thanks also to P. Belin, M. Besson, M. Brett, J. Duncan, W. Freiwald, A. Martinez, J. Iglesias, E. Olivares and P. Valdes-Sosa for comments on the manuscript, and to S. Kastner and M. Arcaro for help with the retinotopic atlas. We also thank three anonymous reviewers for their comments.

This work was supported by the VLIR-UOS project “A Cuban National School of Neurotechnology for Cognitive Aging”, the National Fund for Science and Innovation of Cuba, the National Natural Science Foundation of China (#81861128001) and the ‘111′ Project (B12027) of China.

## Supplementary material

### 1. Figures and tables

**Figure S1.**
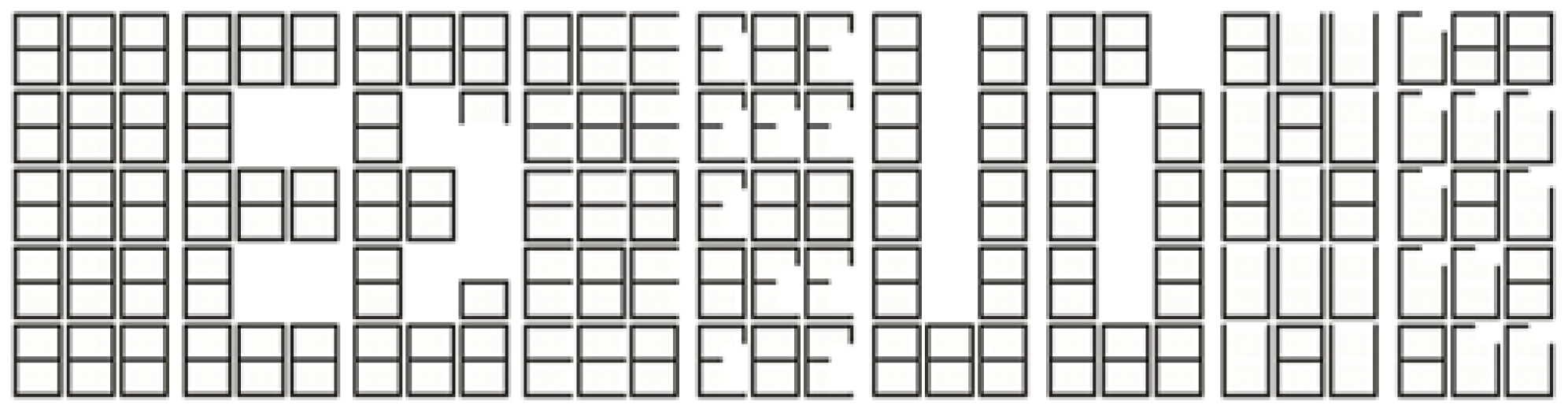
All stimuli used in the main fMRI experiment.

**Figure S2.**
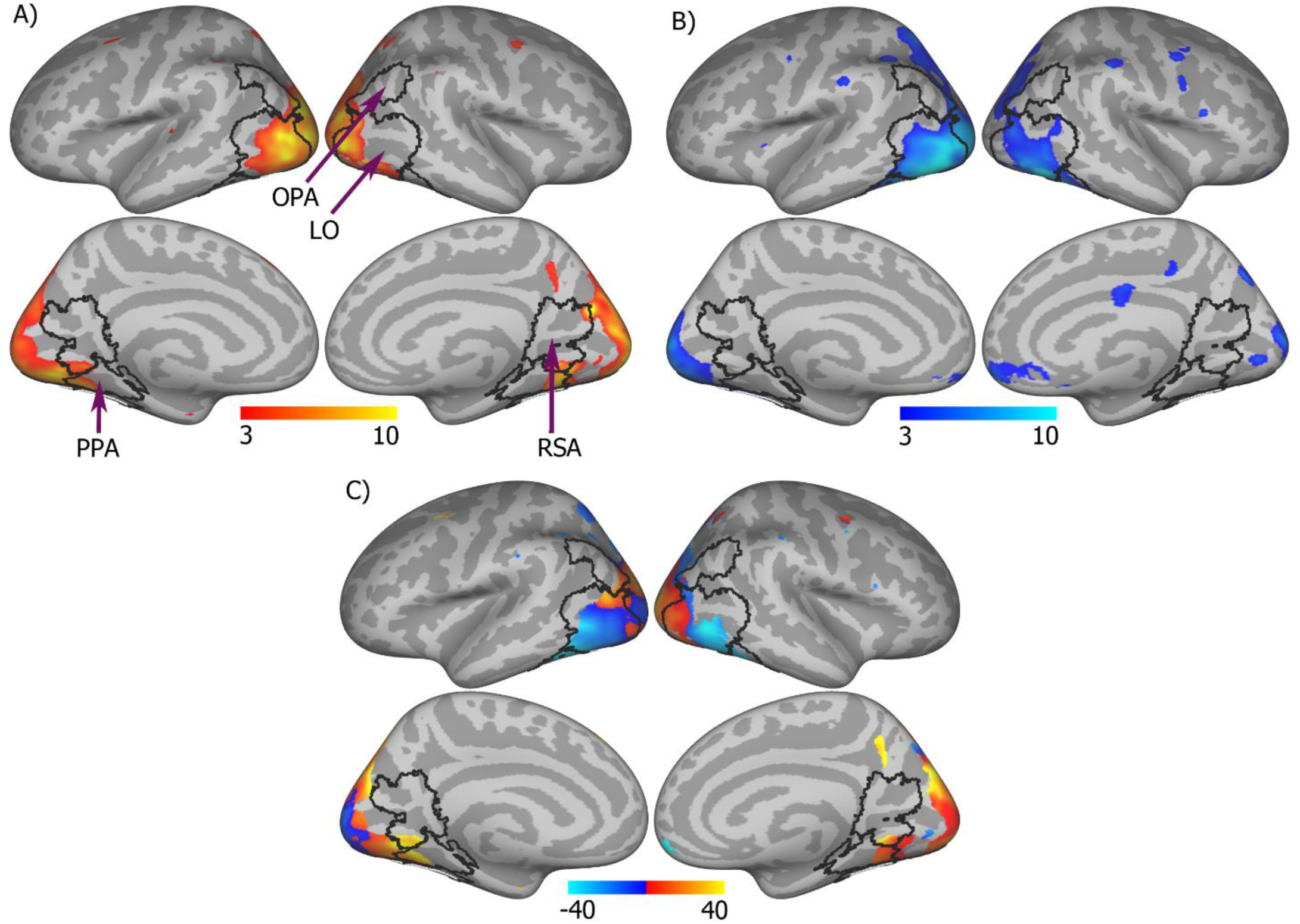
Additional views of searchlight maps for original data: A) Shape-invariant level map. B) Level-invariant shape map. C) IOS map.

**Figure S3.**
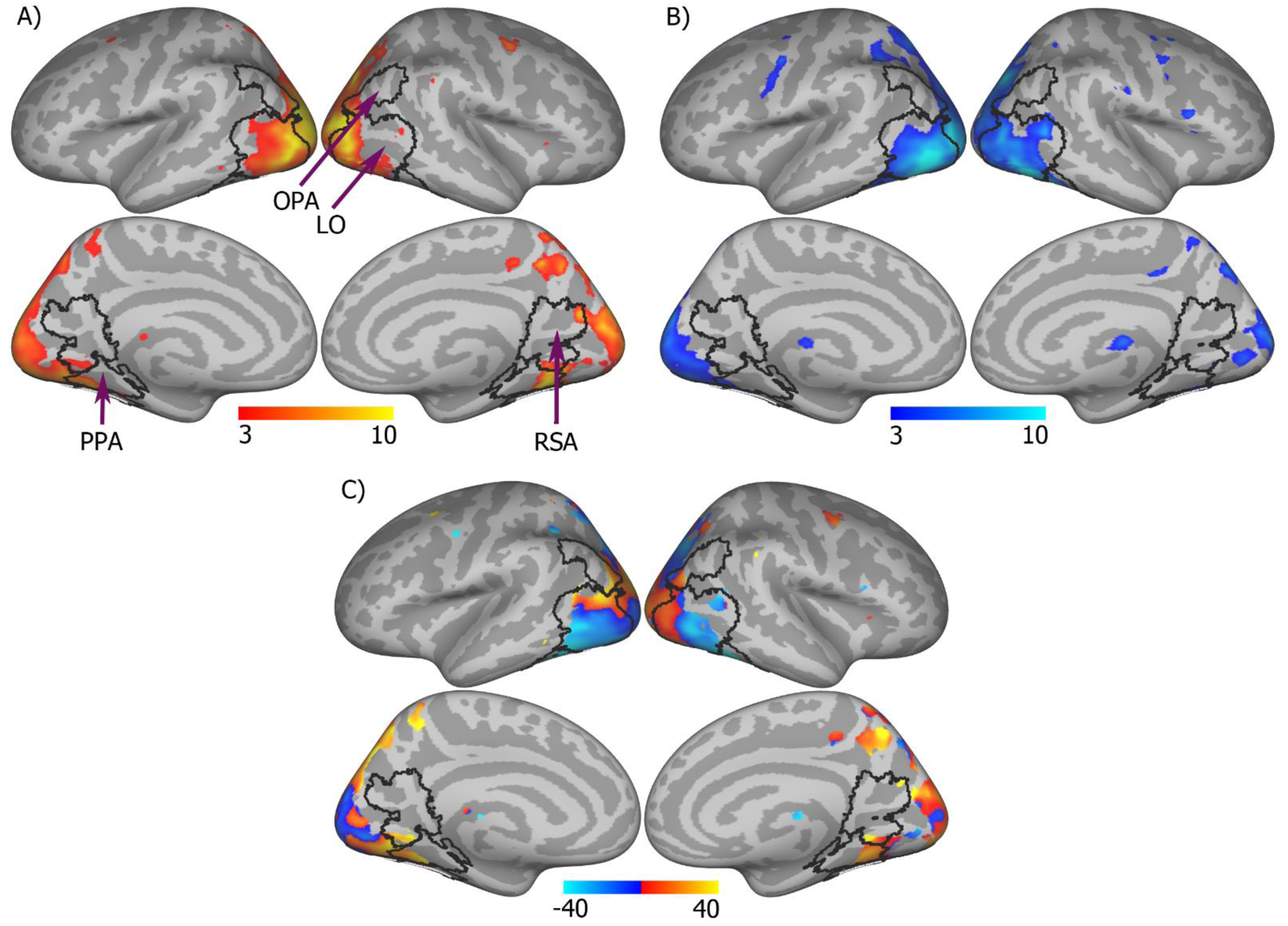
Additional views of searchlight maps for centered data: A) Shape-invariant level map. B) Level-invariant shape map. C) IOS map.

**Figure S4.**
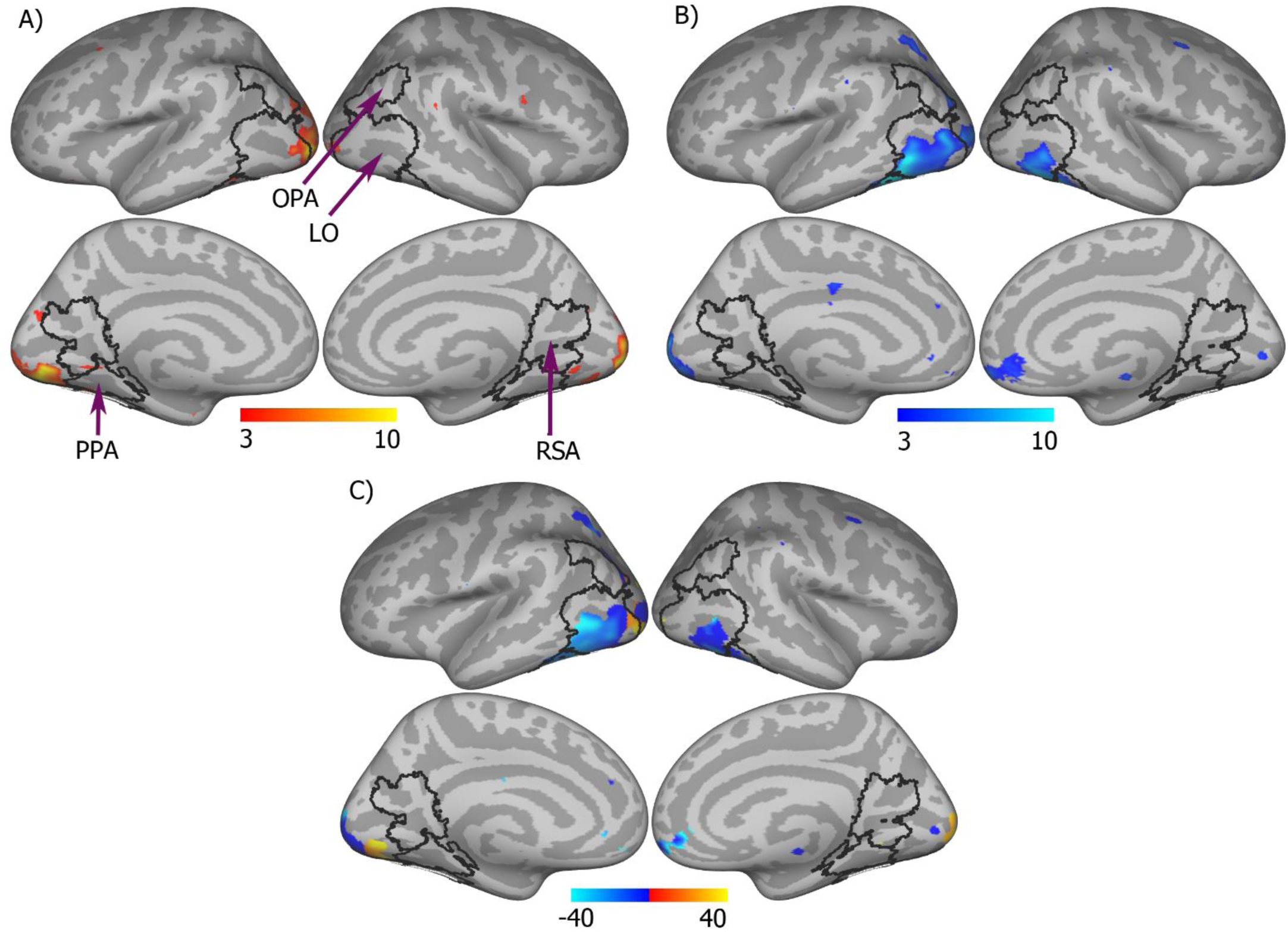
Additional views of searchlight maps for smoothed data: A) Shape-invariant level map. B) Level-invariant shape map. C) IOS map.

**Figure S5.**
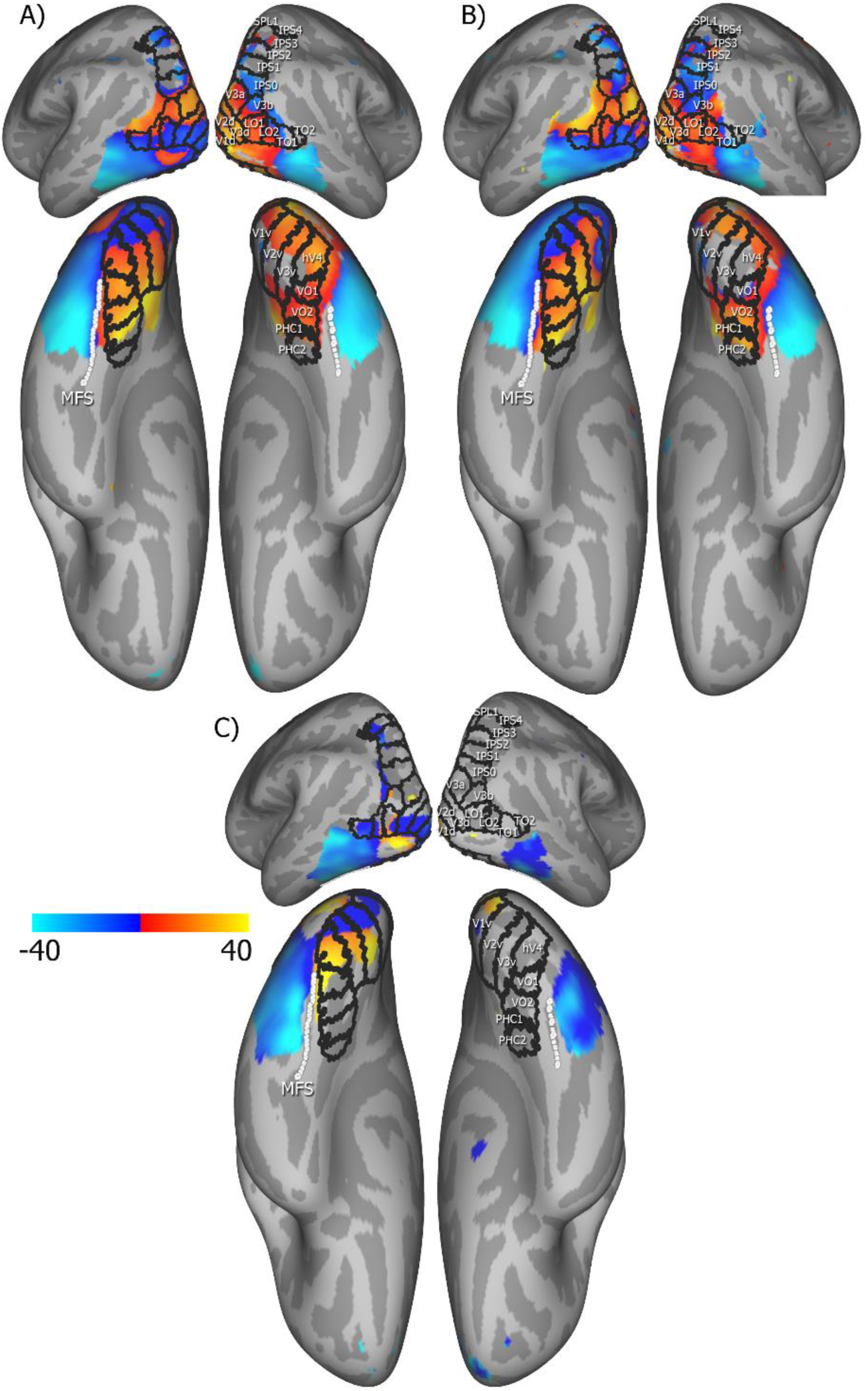
IOS map with superimposed borders of retinotopic areas (Wang atlas). A) Original map. B) Centered map. C) Smoothed map.

**Table S1.** Percentage of informative searchlight centers in each fROI. Percentages larger than 20% are shown in bold.

| fROIs | Percentage (LH) |  |  |  | Percentage (RH) |  |  |  |
| --- | --- | --- | --- | --- | --- | --- | --- | --- |
|  | Houses | Level | Shape | Faces | Houses | Level | Shape | Faces |
| PPA | **58** | 19 | 0 | 0 | **76** | 19 | 0 | 0 |
| RSC | 0 | 9 | 0 | 0 | 15 | 6 | 0 | 0 |
| OPA | 19 | 4 | 0 | 0 | 19 | 0 | 1 | 0 |
| LO | 8 | 5 | **36** | 0 | 9 | 19 | **23** | 8 |
| pFus | **21** | 2 | **41** | **32** | 10 | 0 | **33** | **47** |
| V1v | 1 | 1 | 2 | 0 | 0 | 19 | 4 | 0 |
| V1d | 0 | 6 | 30 | 0 | 0 | **36** | 0 | 0 |
| V2v | 1 | **27** | 0 | 0 | 0 | **43** | 0 | 0 |
| V2d | 2 | 0 | 14 | 0 | 0 | **92** | 0 | 0 |
| V3v | 5 | **40** | 0 | 0 | 7 | **42** | 0 | 0 |
| V3d | 13 | **23** | 9 | 0 | 9 | **93** | 0 | 0 |
| hV4 | 19 | **24** | 0 | 0 | 9 | **78** | 0 | 0 |
| VO1 | **73** | **93** | 0 | 0 | **55** | **46** | 0 | 0 |
| VO2 | **91** | **98** | 0 | 0 | **75** | **74** | 0 | 0 |
| PHC1 | **97** | **54** | 0 | 0 | **100** | **87** | 0 | 0 |
| PHC2 | **100** | 4 | 0 | 0 | **100** | 17 | 0 | 0 |
| MST | 0 | 0 | 0 | 0 | 0 | 0 | 0 | 0 |
| hMT | 3 | 1 | 16 | 0 | 9 | 0 | 2 | 14 |
| LO2 | **89** | 0 | 0 | 0 | **81** | **23** | 0 | 0 |
| LO1 | **78** | 0 | 0 | 0 | **94** | **71** | 0 | 0 |
| V3b | **100** | 17 | 0 | 0 | **100** | 12 | 8 | 0 |
| V3a | 13 | **96** | 0 | 0 | 15 | **52** | 0 | 0 |
| IPS0 | **28** | 1 | 10 | 0 | **52** | 0 | **23** | 0 |
| IPS1 | 0 | 1 | 11 | 0 | 4 | 0 | 11 | 0 |
| IPS2 | 0 | 7 | 12 | 0 | 0 | 0 | 0 | 0 |
| IPS3 | 0 | 0 | **43** | 0 | 0 | 1 | 0 | 0 |
| IPS4 | 0 | 0 | 0 | 0 | 0 | 0 | 0 | 0 |
| IPS5 | 0 | 0 | 0 | 0 | 0 | 0 | 0 | 0 |
| SPL1 | 0 | 0 | 2 | 0 | 0 | 0 | 0 | 0 |
| FEF | 0 | 0 | 0 | 0 | 0 | 0 | 0 | 0 |

### 2. Comparison of MVPA with face- and scene-selective areas

#### Localizer experiment

Face- and scene- selective areas were localized using a standard paradigm (Epstein and Kanwisher, 1998) in an independent group of participants composed of sixteen Cuban participants, with ages from 25 to 36 years (7 females). All participants had normal (or corrected-to-normal) vision and were right handed except for one case. None had a history of neurological or psychiatric disease. The experimental procedures were previously approved by the ethics committees of the Cuban Center for Neuroscience, and participants gave written informed consent. One scanning run was used, each consisting of 16 blocks of stimuli. The blocks lasted 30 s and each containing 60 stimuli, which in turn were presented during 500 ms. In half of these blocks pictures of faces were presented, and on the other half pictures of houses. Both types of blocks were alternated, separated by 30 s of fixation. All stimuli had the same dimensions (6.2° × 8.25°) and were presented in grayscale on black background. Stimuli were projected on a screen behind the participant’s head, viewed through an angled mirror fixed to the MRI head-coil, and generated using the Cogent Matlab toolbox. The data were acquired with a Siemens MAGNETOM Allegra 3T scanner (Siemens, Erlangen, Germany) at the Brain Mapping Unit of the Cuban Center for Neuroscience (CNEURO) using a single channel receiver head coil.

High-spatial resolution functional images (1.8 × 1.8 × 1.8 mm) were collected using a T2*-weighted echo planar imaging sequence, with 47 axial slices, interleaved, effectively covering the entire brain. The parameters for image acquisition were as follows: TR = 3s, TE = 30 ms, flip angle = 90 °, acquisition matrix = 128 × 120. For each subject, 340 volumes were collected. A 176 slice anatomical T1-weighted image was obtained with the following parameters: voxel size = 1 × 1 × 1 mm; TR = 1940 ms; TE = 3.93 ms; acquisition matrix = 256 × 256; and flip angle = 9.

#### Analysis of the localizer data

The localizer data was analyzed as the main experiment up to the massive GLM to estimate the beta values, except that the data was smoothed on the surface with an 8 mm Gaussian kernel. Then, the contrasts of the face>house, face>fixation-point, and house>fixation-point were calculated. The latter two contrasts were entered into a group t-test vs zero and thresholded with an FDR at q=0.05 (based on an empirical null distribution), and were also expressed as an IOS values (see Fig. S6B). This face/house-IOS was calculated as the polar angle between the mean face>fixation-point, and house>fixation-point contras for all nodes in which at least one of the t-tests was significant.

To further test the relationship between level/shape information with the category-specific regions, we performed a more detailed analysis limited to VTC region using the face>house contrast. These contrast values were averaged in each individual within the level/shape masks created with the IOS, separately for the two hemispheres. The averaged contrast values were then submitted to a repeated measures-ANOVA using Category (face/scene) and Hemisphere as main effects. Subsequently, the Pearson correlation coefficient was calculated between the average IOS values and the results of t-tests for the face/house contrast. Note that in this analysis the dependent and independent measures were defined in by experiments in separate groups of participants.

#### Characterization of IOS regions using a face/house localizer

The location of shape-invariant level information in scene-selective cortex motivated a further test taking advantage of the fact that houses reliably activate scene-selective cortex. The face/house-IOS map showed remarkable analogies with the shape/level IOS map. The level/shape divide in VTC across MFS was mirrored by a house/face separation. Again the face/house border corresponded better to MFS than the borders of PPA, LO and pFus. In LO the transition from shape at the antero-ventral towards level in the posterior-dorso portion was mirrored by a face/house transition (Fig. S6A).

The ROIs specialized for level had larger responses for houses than for faces, in both the left (t(15)=-7.08, p<0.001) and right hemisphere (t(15)=−4.1, p<0.001). The ROIs for shape had larger responses for faces than for houses in the right hemisphere only (t(15)=4.1, p<0.001) (Fig. S6C). A strong correlation was found (r=−0.64, p<10^−5^) across cortical nodes between the IOS and t-test scores for the face/house contrast (Fig. S6D) in the VTC.

**Figure S6.**
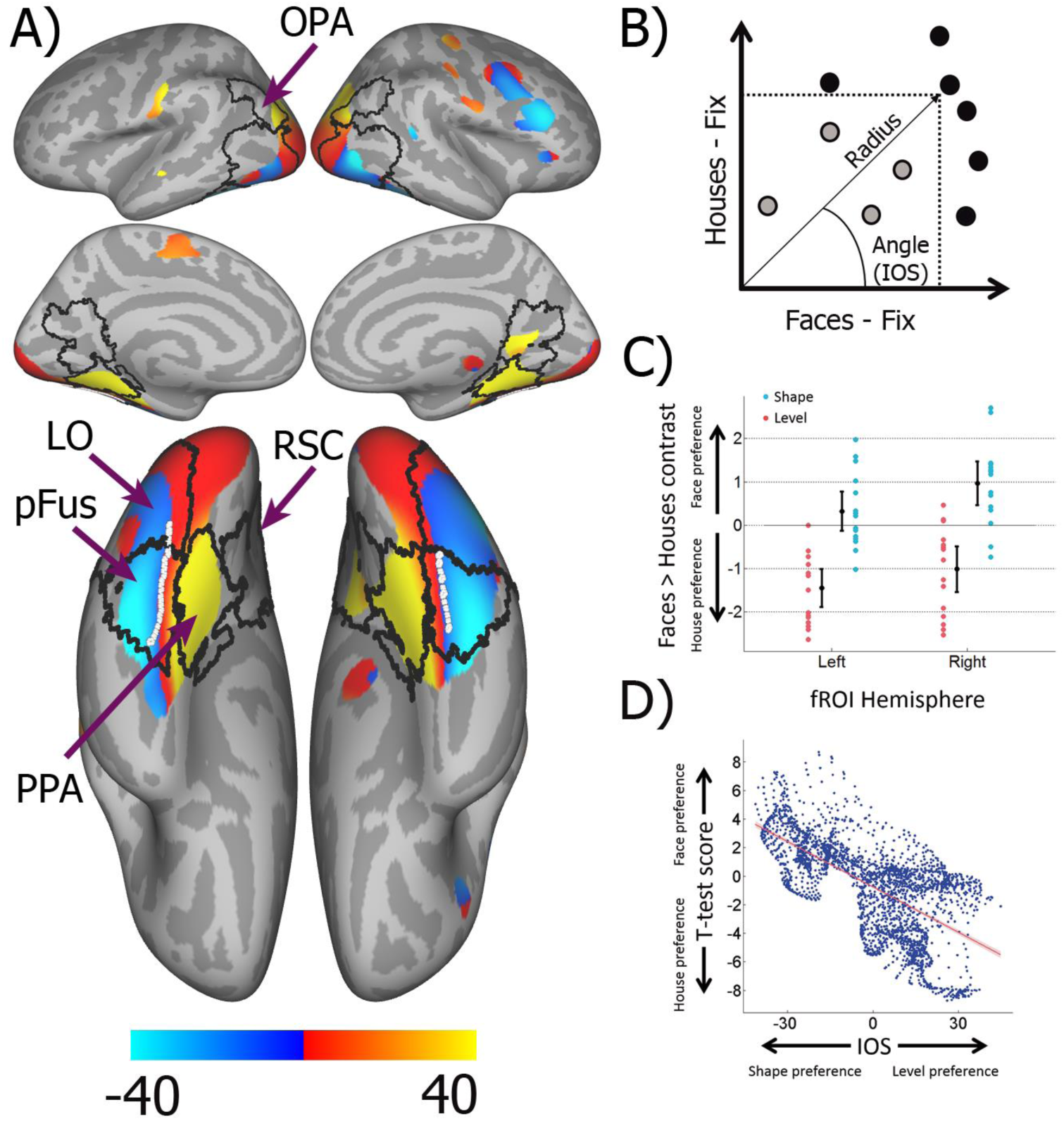
Relationship of regions where level and shape information was invariant with scene- and face-selective areas. A) Group face/house-IOS map. B) Face/house-IOS defined as the polar angle between the contrasts at each node (minus 45 degrees). Dotted lines are FDR thresholds for each t-test. Only nodes significant in at least one t-test were considered (drawn as black circles in this toy example). ROI name conventions as in Fig. 2. C) Scatterplots of mean value of face-house contrast within masks of cortex specialized for level (red dots) and for shape (blue dots) in each individual, separated by hemisphere. Black dots and whiskers represent means and standard errors of means respectively for each ROI. D) t-test scores of the face-house contrast as a function of IOS plotted for cortical nodes in VTC (excluding V1 and V2).

